# In-line swimming dynamics revealed by fish interacting with a robotic mechanism

**DOI:** 10.1101/2022.07.14.500016

**Authors:** Robin Thandiackal, George V. Lauder

## Abstract

Schooling in fish is linked to a number of factors such as increased foraging success, predator avoidance, and social interactions. In addition, a prevailing hypothesis is that swimming in groups provides energetic benefits through hydrodynamic interactions. Thrust wakes are frequently occurring flow structures in fish schools as they are shed behind swimming fish. Despite increased flow speeds in these wakes, recent modelling work has suggested that swimming directly in-line behind an individual may lead to increased efficiency. However, no data are available on live fish interacting with thrust wakes. Here we designed a controlled experiment in which brook trout, *Salvelinus fontinalis*, interact with thrust wakes generated by a robotic mechanism that produces a fish-like wake. We show that trout swim in thrust wakes, reduce their tail-beat frequencies, and synchronize with the robotic flapping mechanism. Our flow and pressure field analysis revealed that the trout are interacting with oncoming vortices and that they exhibit reduced pressure drag at the head compared to swimming in isolation. Together, these experiments suggest that trout swim energetically more efficiently in thrust wakes and support the hypothesis that swimming in the wake of one another is an advantageous strategy to save energy in a school.

## INTRODUCTION

Individuals in fish schools have long been hypothesized to benefit from hydrodynamic advantages associated with swimming near other conspecifics (1–4). Recent work supports this hypothesis on the basis of experiments where schooling fish exhibit reduced tail-beat frequencies relative to solitary individuals which suggests decreased energy consumption by the group as a whole (5, 6). A number of specific mechanisms have been proposed and investigated to show how corresponding hydrodynamic effects could contribute to reduced energy demands in schools (**Fig. 1**). The phalanx or soldier formation describes fish swimming side-by-side, parallel to each other (**Fig. 1A**), and fish in this position are expected to benefit from the channeling/wall effect and simulation studies (7, 8) have shown increased efficiency for this formation. And Ashraf et al. (5) linked the phalanx formation to reduced energy consumption in red nose tetras swimming in a school. Another beneficial interaction can occur when two fish swim in close proximity to one another (**Fig. 1B**). Here, the leading swimmer is thought to experience increased thrust because of the additional effective added mass at the tail trailing edge due to blockage of water by the trailing swimmer behind. Simulations on pitching foils (9, 10) have confirmed this effect and show increased overall hydrodynamic efficiency for the two-body system of leading and trailing swimmers. Measurements of reduced tail-beat frequencies of fish swimming at the front of schools of grey mullet compared to swimming in isolation further support these findings (6).

**Fig. 1:**
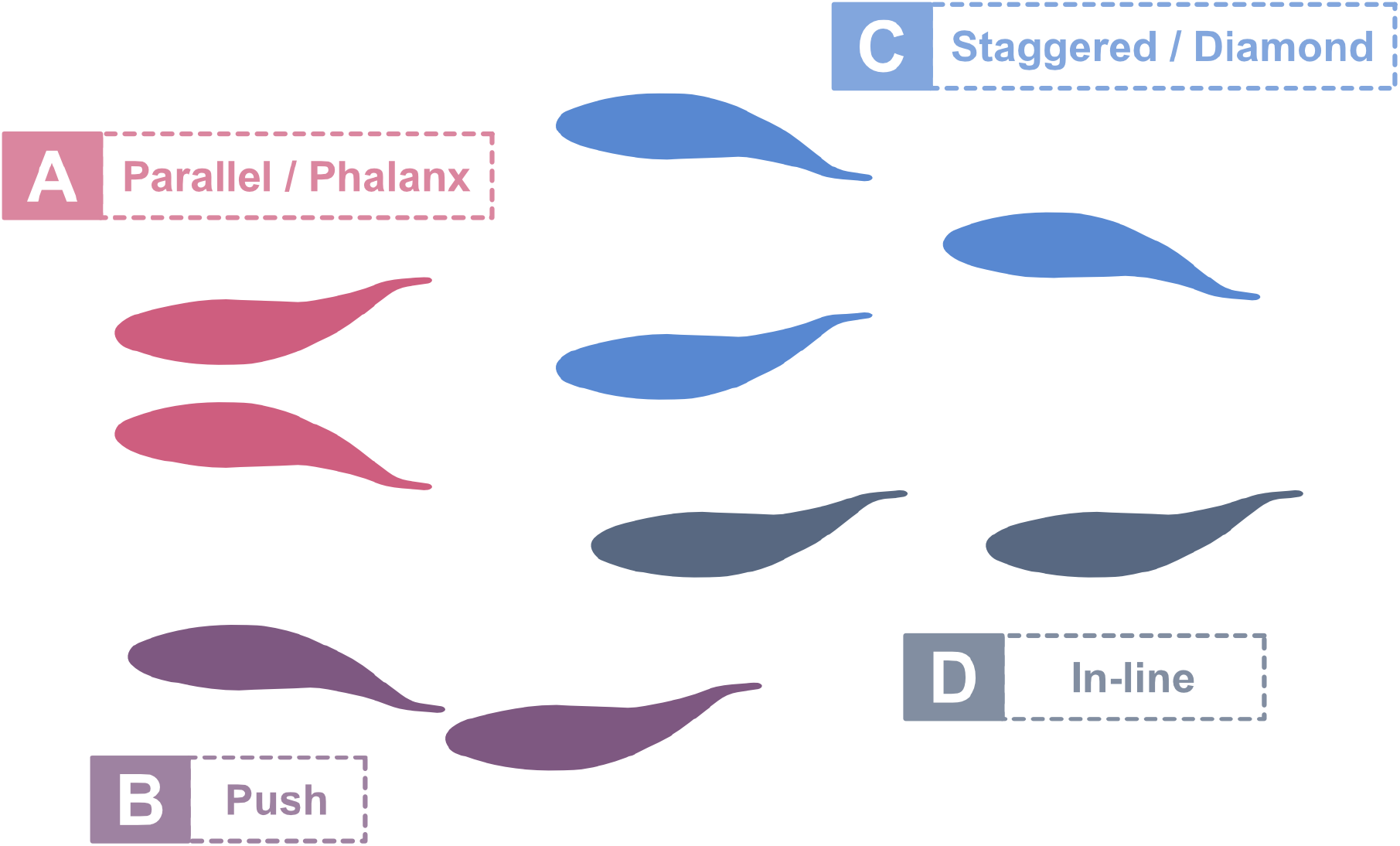
Schooling positions with hydrodynamic benefits. **(A)** Swimming side-by-side can increase thrust and efficiency by making use of the channeling effect (5, 7). **(B)** Leading swimmers benefit from higher thrust production due to increased effective added mass at their trailing edge stemming from the blockage of the water in close proximity to trailing swimmers (9, 10). **(C)** Trailing fish face reduced oncoming flows between two leading fish when swimming in a diamond formation (4). **(D)** Leading-edge suction provides propulsive thrust for a fish in a trailing position (9, 17, 19).

A third commonly proposed schooling arrangement is the diamond or staggered pattern (**Fig. 1C**) first suggested by Weihs (4). The value of swimming in this formation is due to the nature of thrust wake vortical structures generated behind swimming fish. Fish thrust wakes are characterized by both a vortex street of alternating orientation, and an increased average flow speed compared to the free stream (11–14). Weihs hypothesized that fish directly behind another *“would experience a higher relative velocity and would have to exert extra energy”* and suggested that the most efficient swimming position lies midway between two preceding fish (**Fig. 1C**) resulting in a diamond formation. A fish swimming in this diamond formation encounters flow conditions resembling a von Kármán drag wake, similar to the one shed by a cylinder under sufficiently high flow speeds. Liao et al. (15) explored this scenario in trout and found reduced muscle activity for fish swimming in a drag wake, and direct measurements of energy consumption confirm that fish experience reduced energetic costs when in a drag wake (16).

In contrast to Weihs’ argument that in-line fish positions are disadvantageous (**Fig. 1D**), some recent work suggests that swimming in tandem provides hydrodynamic advantages. Simulations (8, 17), flapping foil experiments (18, 19), and robot experiments (9) indicate increased thrust production and efficiency when a fish or flapping foil swims in a thrust wake. The fluid dynamic benefits to the follower occur because the swimmer in the thrust wake experiences the oncoming flow at its leading-edge with an oscillating angle of attack and is subject to lift forces that have components in forward direction. Maertens et. al (17) argue that a downstream swimmer *“can reduce its drag by consistently turning its head in a manner that employs the oncoming vortex flow to increase the transverse velocity across the head”*. As a result, the pressure drag at the head can be decreased substantially and result in increased efficiency.

Although recent modeling work suggests advantages for in-line swimming, experimental data on live fish exploiting these conditions is lacking. Do live fish actually take positions in a thrust wake when free to swim at any location in flow? When fish swim directly behind another, do they alter their swimming kinematics and is there evidence for a reduction of swimming cost even when in a thrust wake with accelerated mean flow? Here we explore how fish interact with thrust wakes in a controlled experimental setting. We chose trout (brook trout, *Salvelinus fontinalis*) for our investigation as this species swims against oncoming currents in their natural habitat and is known to sense and take advantage of flow structures that can reduce energy use (20, 21). In our approach we emulate the thrust wakes from leading swimmers using an actuated flapping foil that serves as the artificial counterpart of a fish tail-fin to generate accelerated flows with similar hydrodynamic characteristics to those of live fish (22, 23). By carefully choosing the robotic flapping motion, we generated fish-like thrust wakes and introduced trout to these conditions. We found that trout swim in-line with the flapping foil (**Supplementary movie 1 and 2**) and reduce their tail-beat frequencies compared to swimming at the same effective flow speeds under free-stream conditions. Further analyses employing particle image velocimetry revealed that individuals interact directly with oncoming thrust wake vortices. Finally, our pressure field computations showed reduced average pressures at the leading-edge suggesting reduced pressure drag and reduced swimming costs. These findings support the hypothesis that fish can reduce swimming costs under in-line swimming conditions and help explain why in-line swimming is common in schools of fish.

## RESULTS

### Reduced frequency and synchronization with a flapping foil

Artificial thrust wakes were generated in a recirculating flow tank using an actuated flapping foil with 2 degrees of freedom which enabled side-to-side movement as well as rotation (**Materials and Methods: Flapping foil, Fig. S1**). The motion of the foil together with the flow speed (*St* = 0.267) were chosen such that the Strouhal number falls in the typical range of 0.2-0.4 for swimming fish (24). The thrust wake generated by the flapping foil is characterized by a reverse Kármán vortex street and increased flow speeds in the wake (**Fig. S2**) comparable to the wakes generated by swimming trout (11).

We used a paired experimental design and had the same individuals swim under two conditions: in a flow tank with (1) an actuated flapping foil generating a thrust wake, and (2) under control free stream conditions with the foil in a stationary position. In both conditions the flow was fixed at the same speed, and thus permitted a fair comparison of the corresponding swimming patterns. In addition, we carried out the same experiments 2.5 months apart, which allowed us to investigate how differences in body size affect the behavior under the different conditions as fish were larger in total length after this growth period. We captured the swimming kinematics using high-speed video recordings from the ventral perspective and extracted body midlines (**Materials and Methods: Experimental setup and Kinematic Analysis**).

We found that trout from both size groups significantly reduced their tail-beat frequencies when they were exposed to thrust wakes (**Fig. 2A, supplementary movie 2**). Smaller fish showed a decrease of 28.3 %, and larger fish showed a decrease of 14.7 % in the mean frequency. This suggests that fish maintained their position in the thrust wake by beating their tails less often than when they swam at the same ground speed in free-stream flow. These experiments further showed that fish were synchronizing their tail-beat frequency to that of the flapping foil. For both smaller (mean±s.d.: 2.15 ± 0.29 Hz) and larger (1.96 ± 0.04 Hz) fish the swimming frequency approached the 2 Hz flapping foil motion when swimming in the foil thrust wake.

**Fig. 2:**
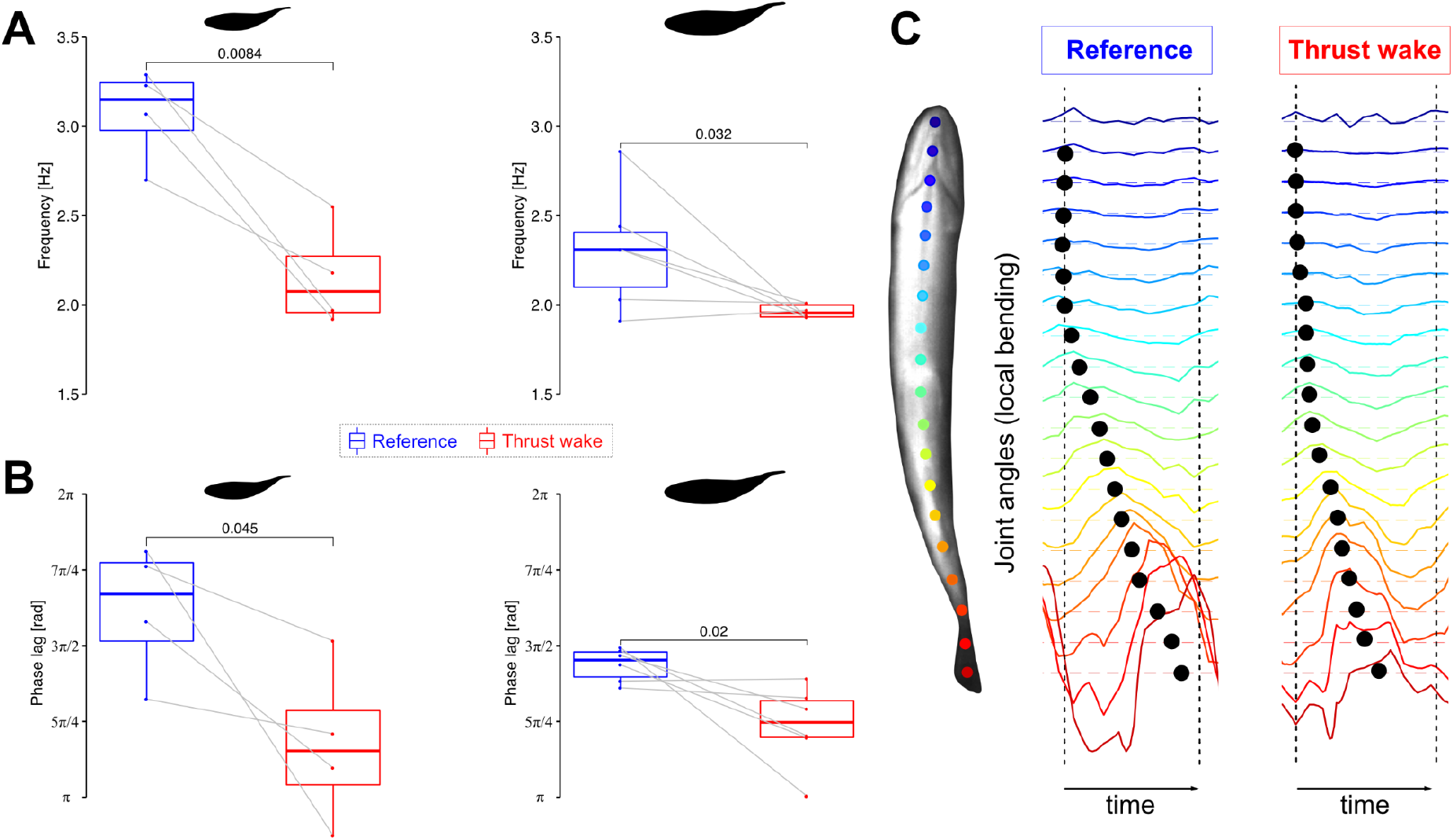
Body kinematics in thrust wakes. **(A)** Reduced tail-beat frequencies and **(B)** reduced overall phase lags for small (n=4) and large (n=6) trout swimming in the thrust wake compared to steady swimming at the same flow tank speed. **(C)** Illustration of the bending pattern by means of joint angles (rainbow colored lines) along the body. Black markers indicate the bending phase.

We analyzed body bending kinematics and identified decreased overall phase lags along the body in the thrust wake in both size groups (**Fig. 2B**) compared to the free stream control condition. As a result, the bending of consecutive body segments was timed closer together (**Fig. 2C**). Smaller overall phase lags also relate to fewer waves along the body. We did not find any significant differences in body amplitude between fish that swam in thrust wakes and in the free stream (**Fig. S3**).

### How do fish interact with the flow in thrust wakes?

Kinematic analysis of fish swimming in thrust wakes indicates a frequency synchronization with the flapping foil. To investigate flow dynamics and how the thrust wake generated by the foil interacts with the bending fish body we employed particle image velocimetry (**Materials and Methods: Setup to capture flow dynamics, supplementary movie 3**) to visualize flow structures in the thrust wake during fish swimming trials. Analysis of flapper wake velocity fields show that trout in the thrust wakes interact with oncoming vortices that are shed from the flapping foil and time their movements accordingly. We identified two scenarios that we call double-sided and single-sided vortex interaction. Double-sided vortex interaction (**Fig. 3A-F, supplementary movie 4**) is characterized by an initial vortex interception which splits the vortex in two parts. One part of the vortex stays attached and “rolls’’ downstream along the body, whereas the other part is shed laterally and moves away from the body. In this situation, the trout body alternates between clockwise and counter-clockwise vortices that are intercepted, stay attached and roll along corresponding alternate sides of the body. Fish are thus able to “catch vortices on both sides of the body”.

**Fig. 3:**
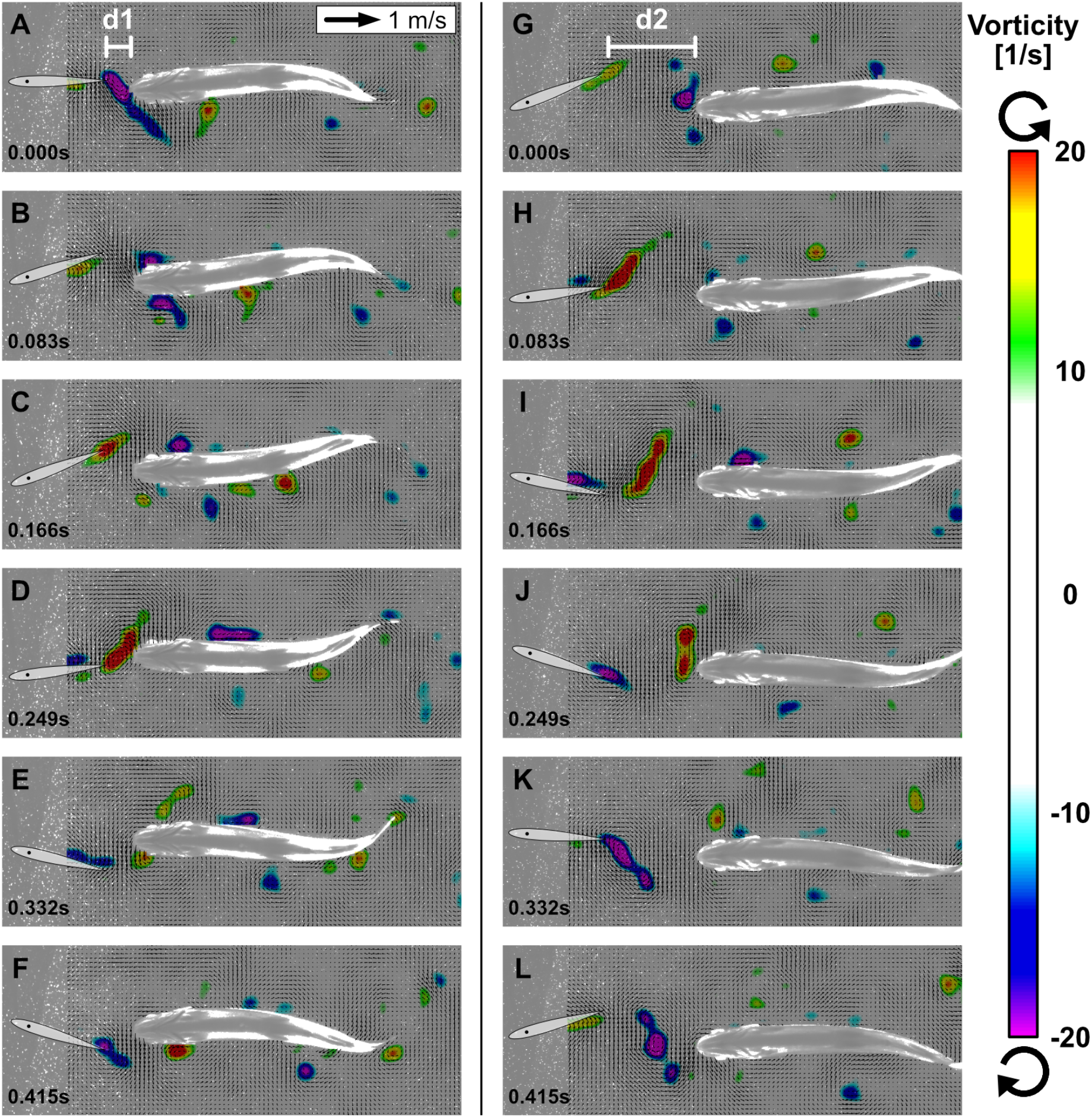
Interactions between fish and vortices. Two representative sequences over one swimming cycle with ventral view of trout station-holding in the thrust wake near the foil at distance d1 with double-sided vortex interactions **(A-F)** and located more downstream at d2 with single-sided vortex interactions **(G-L)**. Oncoming vortices from the flapping foil are intercepted by trout in the wake. The vortices stay attached on one side depending on their orientation and “roll” downstream along the body (velocity fields shown after subtraction of mean flow speed).

Single-sided vortex interactions (**Fig. 3G-L**) undergo the same process of “catching vortices”; however only for one of the two differently oriented vortex types that are shed from the foil. Consequently, the intercepted vortex stays attached and “rolls” only along one side of the body. The single-sided vortex interactions are related to a slight lateral offset of fish position with respect to the center line around which the foil is oscillating, whereas double-sided vortex interactions occur for fish swimming directly in the center line. We also note that vortex interactions occurred at different distances with respect to the foil (d1 and d2 in **Fig. 3**). The consistent interaction pattern between the trout body and oncoming vortices indicates that these fish are synchronizing their movements with respect to the flapping foil and the corresponding vortices shed into the wake.

### Decreased head pressure indicates reduced energy requirements

A large part of the drag on a swimming fish at Reynolds numbers greater than 5000 is caused by drag forces at the anterior portion of the body that faces the oncoming flow (25, 26). Total body pressure drag on a swimming streamlined fish like trout is mainly determined by the pressure acting on the head (*F = p*. *S*, F: drag force, p: pressure, S: surface area). Therefore, to estimate the effect of swimming in a thrust wake on drag we compared pressure fields of fish swimming in the free stream to thrust wake conditions.

We derived the pressure fields at the anterior part of the fish body based on velocity field changes as proposed by Dabiri et al. (27) (**Materials and Methods: Pressure field computation**). The computed pressure fields revealed reduced average head pressures in the thrust wake (**Fig. 4A, B, Fig. S4**). We found the strongest decreases (46% and 86% decrease compared to free-stream swimming) for fish swimming close to the foil and exploiting double-sided vortex interactions. Fish swimming further away from the foil and exhibiting single-sided vortex interactions also showed reduced average head pressure magnitudes compared to free-stream swimming (45% decrease). Here we found an asymmetric average pressure pattern with higher average pressures at the side closer to the centerline of the foil oscillation. The other side of the head experienced smaller average pressures.

**Fig. 4:**
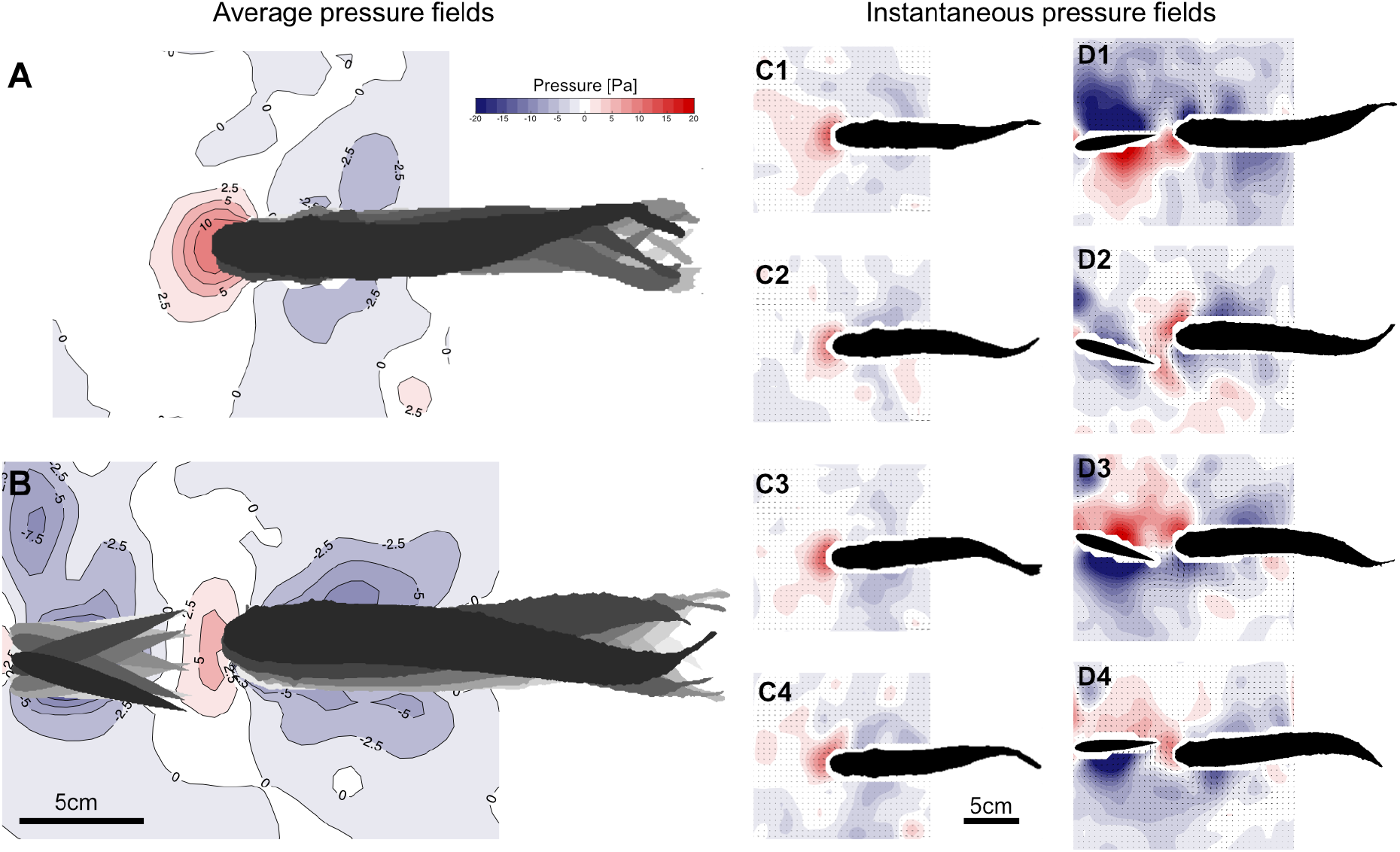
Reduced head pressure in the thrust wake. Average pressure fields of a trout swimming in free-stream flow **(A)** and in the thrust wake of a flapping foil **(B)** show reduced positive pressures (46% decrease) around the head despite increased oncoming flow. Consistent instantaneous positive pressures over time are present under free-stream flow conditions (**C1-C4**). Corresponding instantaneous pressure fields display alternating positive and negative pressures around the head in the thrust wake over time (**D1-D4**).

To understand how the average head pressures in the thrust wake were reduced despite faster oncoming flows caused by the flapping foil, we analyzed the instantaneous pressure fields (**Fig. 4 D1-D4**). Here, it becomes evident that the flapping foil induces oscillating negative and positive pressure zones around the head. The negative pressure (suction) zones cause forward thrust forces, whereas the positive pressures contribute to drag. On average, this reduces overall head drag as the positive pressure magnitudes are comparable to free-stream swimming but mean head pressure is reduced by the occurrence of negative head pressures for part of the cycle. This pressure analysis indicates that the drag of fish swimming in thrust wakes is reduced compared to free-stream swimming, and therefore supports the hypothesis of decreased energy used to hold station in thrust wakes with accelerated mean flow.

## DISCUSSION

In schools of swimming fishes there are a number of different hydrodynamic effects that can be exploited to save energy by individual fish in various positions (**Fig. 1**). Previous work has demonstrated benefits for swimming side-by-side (phalanx configuration), pushing off near followers, and forming diamond patterns (5, 9, 16). But fish in schools often assume an in-line configuration with one fish swimming directly behind another (**Supplementary movie 5**). The benefits, if any, of swimming in this tandem swimming mode have been the subject of some debate. Some authors (4, 28) suggest that swimming in tandem is not an energetically favorable configuration due to the accelerated wake flows generated by the fish in front. Other, primarily computational studies, have suggested that a trailing streamlined shape could in fact experience reduced energetic cost due to leading edge suction resulting from an oscillating flow impinging on the head or leading edge of the trailing fish or foil (9, 17, 19). To date, however, no experimental study has demonstrated that live fish will voluntarily swim in a thrust wake and that reduced swimming cost could result from such a position. With these experiments we document that trout indeed perform volitional in-line swimming with their body located within the accelerated flow region, and our analysis suggests that they can save energy under these conditions.

### Comparisons to drag wake swimming and differences from drafting

A drag wake in the context of fish swimming and schooling is characterized by a von Kármán vortex street between two thrust wakes (e.g., shed by two fish swimming parallel to each other). Drag wakes can be emulated behind cylinders when they are exposed to sufficiently high flow speeds. It is important to note that the average flow speed in a drag wake is inherently slower than in the free stream. This is highlighted in experiments that demonstrate a dead fish propelling itself forward in the wake of a cylinder (29). Intuitively, we can draw an analogy of a cyclist drafting behind another cyclist where the individual behind experiences reduced energy consumption while maintaining the same speed. This situation is an example of a drag wake and the reduced costs that ensue from moving in that reduced velocity zone: trailing cyclists benefit from the reduced relative oncoming flow which results in reduced aerodynamic drag.

The dynamics of thrust wakes, however, differ from drag wakes. Vortex orientations are reversed compared to the drag wake (termed a reverse Kármán vortex street), and, notably, thrust wakes are characterized by a higher average flow speed than in the free stream. For swimming fish, this is a consequence of tail fin movement which actively generates thrust so that individual fish located behind this thrust wake experiences higher than free stream mean flow velocities. Given this increased oncoming flow speed it is surprising that fish choose to swim in thrust wakes. If we return to our example of cyclists, it would correspond to a (fictional) case where a leading cyclist would have a propeller attached to their bicycle that generates additional thrust. The trailing cyclist would face an increased oncoming flow and experience increased aerodynamic drag. An in-line, tandem, formation might be expected to be disadvantageous in this case.

How does this situation differ from fish swimming in a thrust wake of a conspecific? A key difference is the undulatory characteristic of thrust wakes that are produced by swimming fish. A trailing fish faces the oncoming flow at its head with an oscillating angle of attack, and (unlike the trailing cyclist) the trailing fish oscillates its head during swimming (30) further enhancing the time-dependent variation in flow in the head region. Our analysis showed that as a result of the oscillatory wake impinging on the fish head the pressure distribution in the head region is composed of both positive and negative pressures, and thus effectively reduces the overall pressure drag. This is in agreement with previous simulation analyses (9, 17) and makes in-line swimming an advantageous formation for fish.

### Swimming efficiency in thrust wakes

In our experiments we found that fish in thrust wakes significantly reduced their tail-beat frequency and the frequency, higher when swimming in the free stream, shifted toward the flapping foil frequency. Ultimately, this enabled trout to interact with the oncoming vortices, and, as explained above, to achieve a lower pressure drag. Apart from the reduced hydrodynamic resistance, the lower tail-beat frequencies without significant changes in amplitude also are reflective of reduced metabolic cost (31, 32). The change in phase lags that we observed further suggests a change in the muscle activation pattern along the body. Decreased muscle activation was observed by Liao et al. (15) when trout swim with a Karman gait in a drag wake, and passive dynamics could therefore be additional sources for energy savings that would however need to be confirmed in future experiments. Overall, our data suggests that fish can swim more efficiently in thrust wakes because they maintain the same swimming speed as in the free stream while facing reduced pressure drag and spending less energy by beating their tail less frequently.

These results are in agreement with past simulation and robot studies. Li et al. (33) used two fish robots and recorded the energy consumption in different relative formations. They identified a vortex phase matching mechanism to achieve energy-efficiency, which is similar to our findings of live fish interacting with vortices. Verma et al. (28) explored simulated leader-follower formations in a reinforcement learning framework. A first set of optimizations with a reward function based on a modified Froude efficiency led to formations in which followers *“settled close to the center line of the leader’s wake”* and showed well-coordinated behavior of the follower with the wake. In the same work, Verma et al. (28) concluded in a second set of simulations that swimming in-line with a leader is not associated with energetic benefits for the follower. It is important to note that this conclusion was drawn based on a swimming strategy in which the follower would strictly try to attain an in-line position regardless of energetic considerations. As a consequence, the optimization led to an increased swimming amplitude, which permitted in-line swimming but at a higher energetic cost. We confirm that in-line swimming by itself is not necessarily energetically beneficial. More importantly, efficient swimming requires the correct timing of interactions with the wake. Rather than viewing in-line swimming as a policy we can see in-line positions as favorable conditions to maintain wake synchronization as we found in our experiments of double-sided vortex interactions.

Finally, in-line swimming has been dismissed as a beneficial strategy in the past considering the diverging characteristics of three-dimensional compared versus two-dimensional wakes (28). In such cases it could be argued that the area around the centerline of the wake is composed of quiescent flow and in-line swimming offers no opportunity to interact with vortices. Whereas these diverging wakes are predominantly found in simulation studies at lower Reynolds numbers (28, 34, 35), we found no evidence for bifurcating wake structures behind trout swimming in the free stream, a finding in line with previous analyses of trout wake flow patterns (12, 13). The artificial thrust wake generated using our robotic flapping foil produced a parallel vortex street similar to our observations of the wake in freely-swimming trout. In addition, given the experimental data on single-sided vortex interactions, we hypothesize that energy-efficient thrust wake interactions could also occur in diverging wakes but with a small offset to the centerline.

We know from past studies that there are a number of hydrodynamically beneficial schooling positions (**Fig. 1**). Our results complement this body of work with regard to the frequently occurring situation of fish swimming in thrust wakes that are shed by leading individuals, a condition frequently encountered within fish schools during in-line locomotion. In our controlled experiments we show that trout volitionally swim in thrust wakes and exhibit advantageous flow interactions with incoming vortices suggesting increased energy efficiency for the in-line swimming condition. These results highlight the hydrodynamic complexity of fish schooling and support a view in which individuals in schools have a variety of opportunities to save energy when they swim side-by-side, in drag wakes, and in thrust wakes behind each other.

## MATERIALS AND METHODS

### Animals

We used brook trout, *Salvelinus fontinalis*, and carried out the same experiments 2.5 months apart to investigate size effects as the total lengths of the fish increased after this growth period.

Smaller fish had a body length of 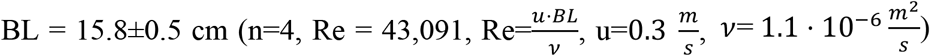 and larger fish had a body length of BL = 19.3±1.0 cm (n=6, Re=52,636). Particle image velocity trials were carried out for larger fish (n=3). Trout were held at a water temperature of 16°C and all experiments were performed in accordance with Harvard animal care and use guidelines, IACUC protocol number 20-03-3 to G.V.L.

### Experimental setup

We carried out all our experiments in a custom flow tank with flow speed control (**Fig. S1, supplementary movie 1**). Fish were able to move freely in a space of 28 cm x 28 cm x 64 cm, where the front and back portions of the swimming section were limited by baffles.

#### Flapping foil

We used a symmetric 3D printed NACA 0012 airfoil (cord: 67 mm; span: 190 mm; thickness: 8.1mm; center of rotation: 48 mm from trailing edge; material: transparent photopolymer (RGD810) from an Connex 500 3D printer) that was actuated in sway and yaw direction (Fig. S1) to mimic the tail-fin portion of swimming fish to induce fish-like thrust wakes. The corresponding movement was parametrized as follows:

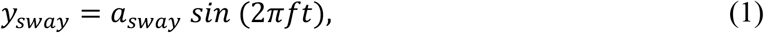

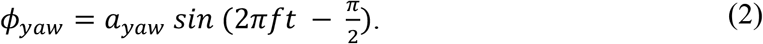

Here, *a*_*sway*_ and *a*_*sway*_ denote the sway and yaw amplitudes, respectively. *f* indicates the flapping frequency and *t* the time in seconds. The two motions are offset by a phase shift of 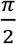, which ensures that maximal yaw is reached whenever the foil crosses zero sway. For the purpose of our experiments *a*_*sway*_ *=* 1cm, *a*_*sway*_ *=* 20° were selected and resulted in a tail-beat amplitude of *A =* 4cm. Together with a frequency of *f =* 2Hz, this resulted in a Strouhal number of 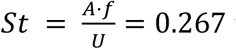 with *U =* 0.3m/s.

#### Fish kinematics

High-speed video was used to capture kinematic variables such as tail-beat frequency, body amplitudes and phase lags during swimming trials. The experiments were carried out in the dark to provide a controlled environment with minimal external distractions. To provide the fish with some sense of visual orientation, a fiber light was installed upstream behind the front baffle in the flow tank. We then used infrared lights (**Fig. S1**), which are outside the visual spectrum of trout, to provide the necessary illumination to capture the scene with high-speed cameras. We took video recordings at 125 frames per second from a ventral and an angled side view.

#### Particle Image Velocimetry

To capture the flow patterns during swimming trials we used particle image velocimetry (PIV) as in our previous work (36–38). For this purpose, we seeded the water in the flow tank with near-neutrally buoyant plastic particles (approx. 50*μ*m mean diameter) and used two lasers to create a light sheet around the swimming fish (**Fig. S1**). Movements were then recorded at 1000 frames per second from a ventral and angled side view (**Supplementary movie 3**). We used the side view to identify the location of fish with respect to the light sheet. Only swimming sequences where the laser light sheet passed through the middle of the swimming fish body were considered in our analysis.

### Kinematic analysis

#### Body midlines

We used a custom MATLAB script to manually track 9 points along the body midline within a given frame. Piecewise cubic spline interpolation was then applied to generate smooth midline curves. We manually tracked the midlines in every 6th frame and linearly interpolated between these frames to obtain midline estimates for all frames that were recorded at 125 frames per second.

#### Frequency estimation

Tail-beat frequencies were determined by averaging the period between maximal lateral tail tip excursions over 3 consecutive swimming cycles for each swimming trial.

#### Phase lag estimation

The phase lag describes how the traveling wave of body bending propagates along the body. To quantify the body bending, we divided the body first into *N =* 20 equal length segments and computed the joint angles *ϕ*_*i*_ between segments. The intersegmental phase lag *Δϕ*_*i*_ was then computed as the time delay between joint angles of consecutive segments as a fraction of the cycle duration *T*. As in (39), we used cross-correlation of the joint angle signals to determine this time delay. Finally, we obtained an estimate of the overall phase lag *Δϕ* by summing up the intersegmental phase lags along the entire body.

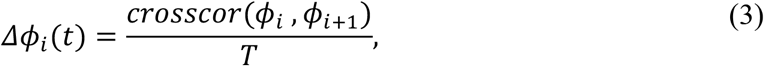

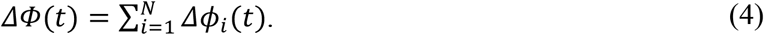

#### Body amplitude estimation

We define the amplitude along the body as the maximal displacement perpendicular to the forward direction of movement. Forward and lateral direction are determined by applying a principal component analysis on the point cloud of all tracked midline points from a given swimming trial. The first principle component (PC) that captures most of the variation represents the forward direction whereas the second PC represents the lateral direction. Based on these directions the lateral displacement at a given midline point in time is then determined as the projection of that point on the lateral direction. Finally, we determine the body amplitude as half of the range of lateral displacements at a given midline point over the duration of the swimming trial.

#### Statistical analysis

To confirm hypothesized decreases in mean frequency and phase lag between the free stream control condition and swimming in the thrust wake, we carried out paired, one-sided Welch t-tests (assuming unequal variance). Significant differences in mean amplitude under these conditions were investigated using paired, two-sided Welch t-tests. P-values are reported in Figures 2 and S3.

### Pressure field computation

Pressure fields were computed based on consecutive velocity fields obtained from particle image velocimetry (**see Experimental setup**). We followed the methodology described in our previous work (40). Velocity fields were computed in DaVis 8.3 (LaVision Inc.) and the pressure fields were obtained using the Queen 2.0 software by Dabiri et al. (27). Corresponding fluid-solid interfaces included both the flapping foil as well as the fish body and were extracted using custom MATLAB and Python scripts.

## DATA AVAILABILITY

Data that support the findings of this study are available on: https://figshare.com/s/b3c687f0b20bf5fdb48a

## ACKNOWLEDGMENTS

This work was supported by the National Science Foundation (Grant number EFRI-830881) and the Office of Naval Research (Grants N00014-18-1-2673, N00014-14-1-0533, and N00014-21-1-2210). We thank members of the Lauder Lab for many helpful discussions about in-line swimming, and Prof. Valentina DiSanto for collaborative research on *Menidia* schooling shown in supplemental movie 5.

## SUPPLEMENTARY FIGURES

**Fig. S1:**
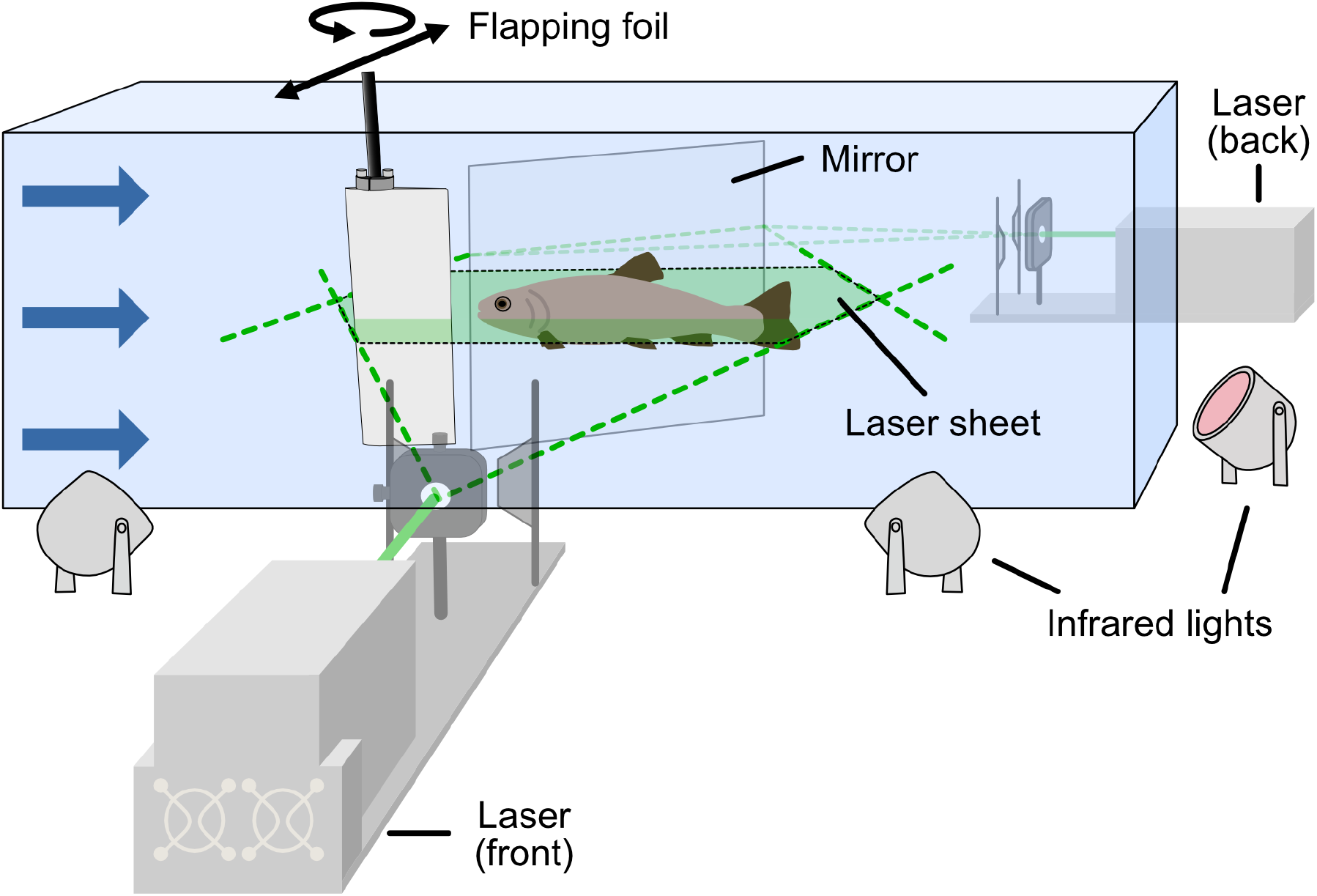
Experimental setup. Flapping foil with 2 degrees of freedom (yaw and sway) generating a fish-like thrust wake in the flow tank. Trout swam in the dark while we captured the kinematics by means of high-speed cameras from a bottom and side view and using infrared lights for illumination. Low light in the tank upstream of the flapping foil allowed fish to orient. In separate experiments, we captured the flow dynamics using particle image velocimetry. We were able to record the entire flow field around the fish by using two lasers (in front and behind) simultaneously.

**Fig. S2:**
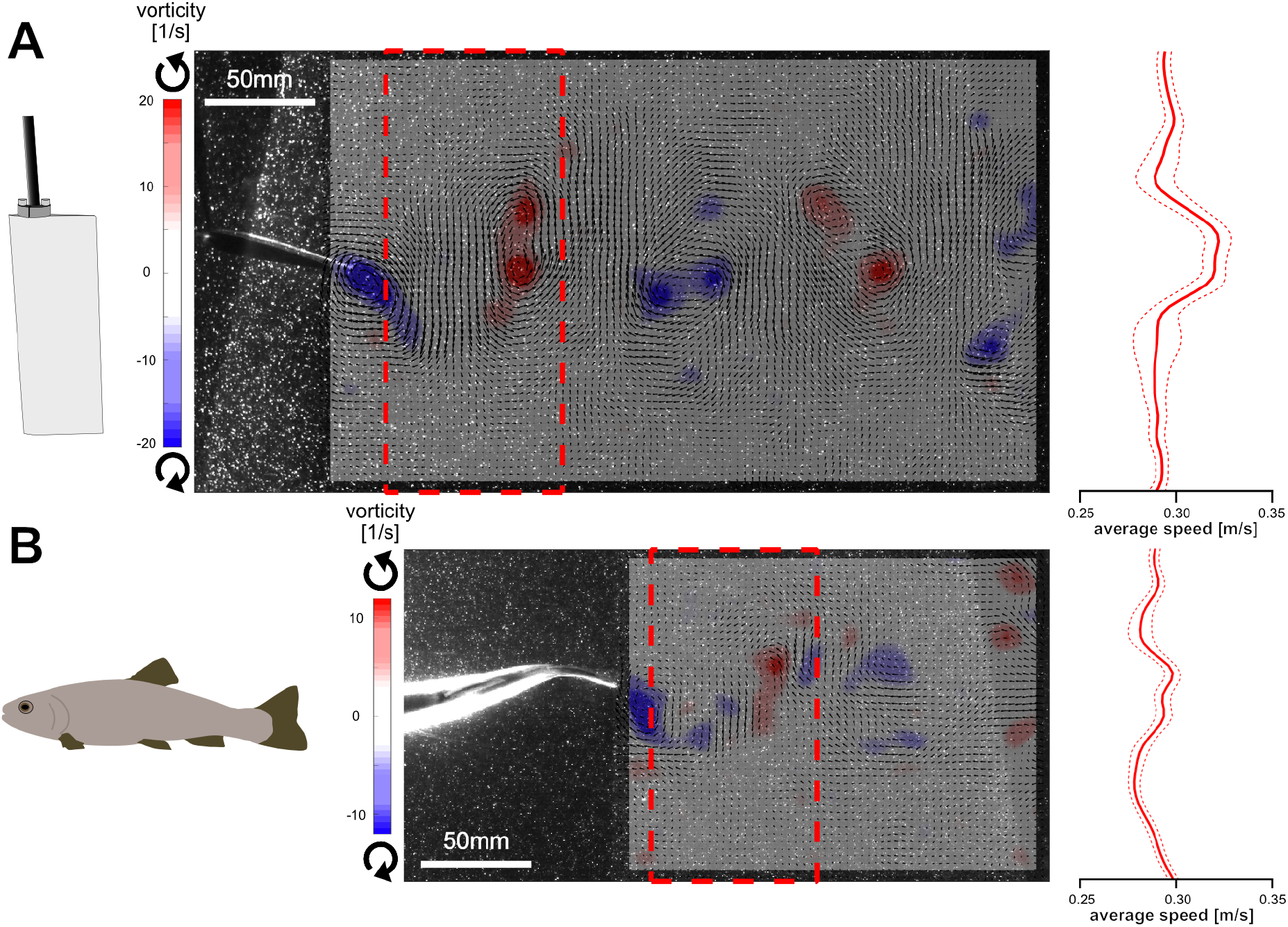
Comparison of flapping foil wake and fish wake. **(A)** Velocity field of the wake behind the flapping foil: red and blue regions indicate counter-clockwise and clockwise vorticity, respectively. The average speed profile of the region within the dashed rectangular region is shown on the right. **(B)** Corresponding plots of the wake behind a steadily swimming trout.

**Fig. S3:**
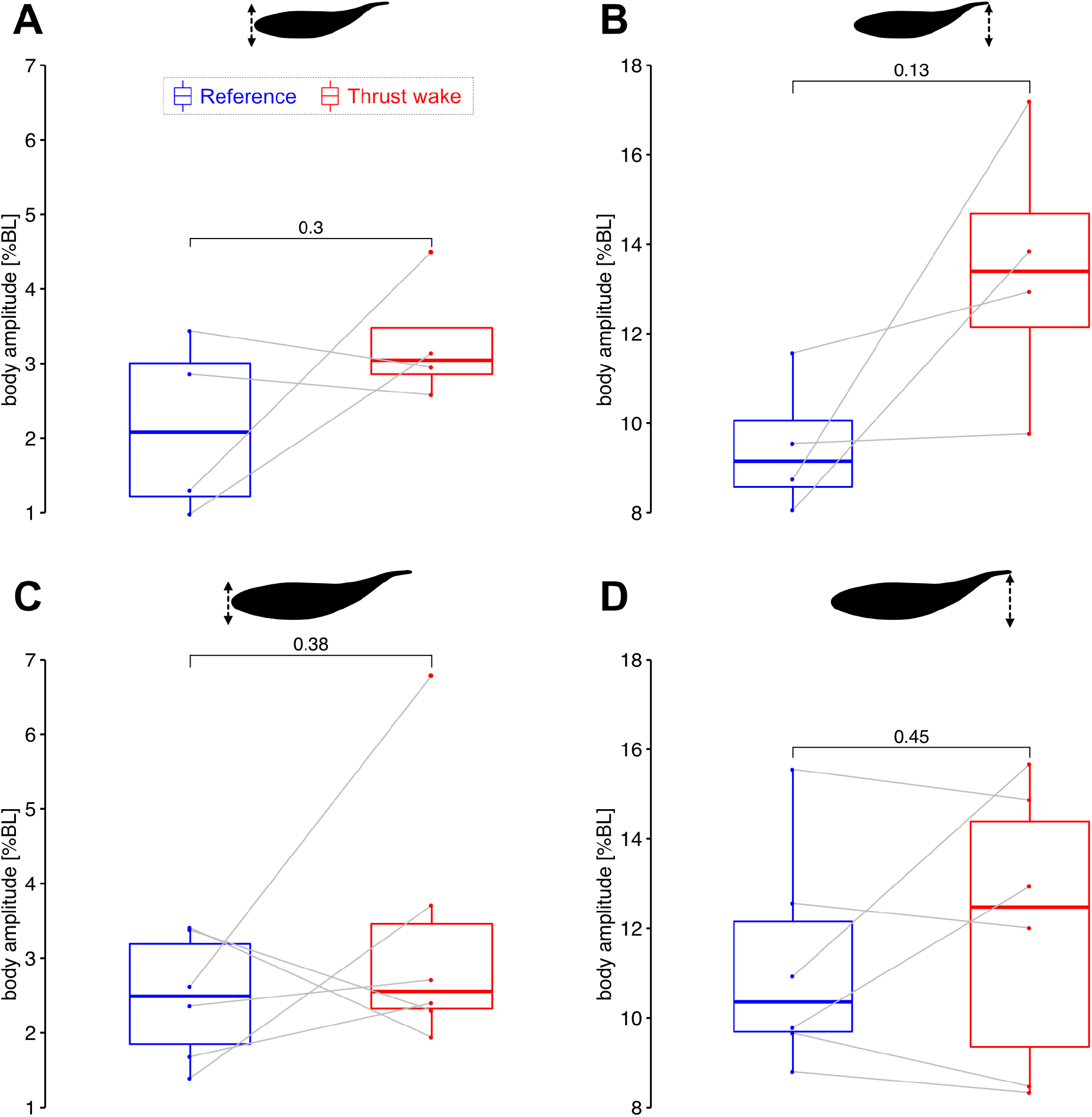
Head and tail amplitudes. Body amplitudes represented as percentage of the body length (BL) for small (n=4) (**A, B**) and large (n=6) (**C, D**) trout. Head (**A, C**) and tail (**B, D**) amplitudes show a slight trend toward increased tail amplitude in the thrust wake but are not significantly different compared to steady swimming.

**Fig. S4:**
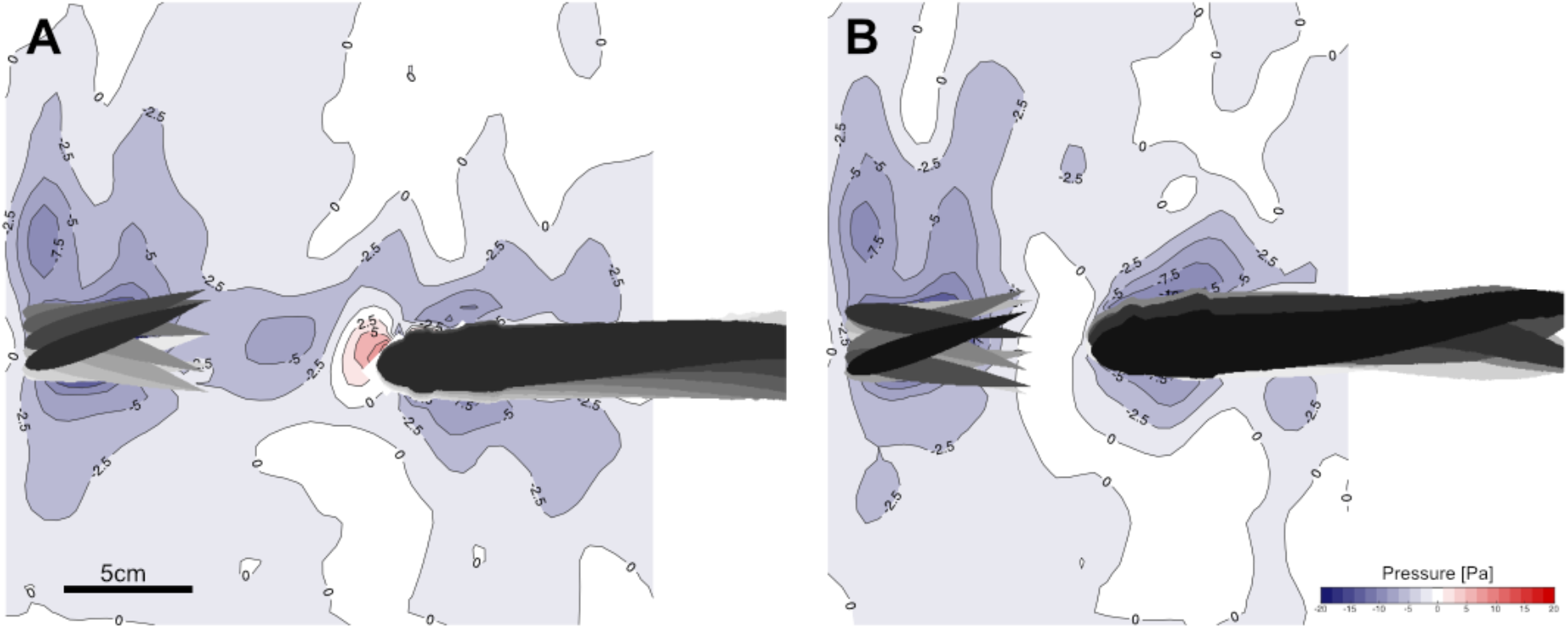
Reduced average head pressures in thrust wakes. **(A)** Trout swimming in the thrust wake exploiting single-sided vortex interactions and 45% decrease in head pressure compared to free-stream swimming. **(B)** Second example of trout exploiting double-sided vortex interactions with 86% decrease in head pressure found.

## SUPPLEMENTARY MOVIES

**Movie 1:** Trout swimming in the thrust wake of a flapping foil (bottom and side views). **Movie 2:** Time lapse of a trout exploring the flow tank with a thrust wake present (bottom view). Over time trout position themselves in the thrust wake and synchronize with the flapping foil.

**Movie 3:** Trout swimming in a laser sheet used for particle image velocimetry (bottom and side views).

**Movie 4:** Visualization of vortex flow structures that interact with a trout swimming in the thrust wake (bottom view).

**Movie 5:** Schooling silversides, *Menidia menidia*, swimming in a flow tank and exhibiting in-line swimming (bottom and side views).

## Notes

### Competing Interest Statement

The authors have declared no competing interest.

## REFERENCES

1. A. D. Becker, H. Masoud, J. W. Newbolt, M. Shelley, L. Ristroph, Hydrodynamic schooling of flapping swimmers. Nat. Commun. 6, 8514 (2015).

2. S. G. Park, H. J. Sung, Hydrodynamics of flexible fins propelled in tandem, diagonal, triangular and diamond configurations. J. Fluid Mech. 840, 154–189 (2018).

3. L. Li, S. Ravi, G. Xie, I. D. Couzin, Using a robotic platform to study the influence of relative tailbeat phase on the energetic costs of side-by-side swimming in fish. Proc. Math. Phys. Eng. Sci. 477, 20200810 (2021).

4. D. Weihs, Hydromechanics of Fish Schooling. Nature 241, 290–291 (1973).

5. I. Ashraf, et al., Simple phalanx pattern leads to energy saving in cohesive fish schooling. Proc. Natl. Acad. Sci. U. S. A. 114, 9599–9604 (2017).

6. S. Marras, et al., Fish swimming in schools save energy regardless of their spatial position. Behav. Ecol. Sociobiol. 69, 219–226 (2015).

7. M. Daghooghi, I. Borazjani, The hydrodynamic advantages of synchronized swimming in a rectangular pattern. Bioinspir. Biomim. 10, 056018 (2015).

8. C. K. Hemelrijk, D. A. P. Reid, H. Hildenbrandt, J. T. Padding, The increased efficiency of fish swimming in a school. Fish Fish 16, 511–521 (2015).

9. M. Saadat, et al., Hydrodynamic advantages of in-line schooling. Bioinspir. Biomim. 16 (2021).

10. Y. Bao, J. J. Tao, Dynamic reactions of a free-pitching foil to the reverse Kármán vortices. Phys. Fluids 26, 031704 (2014).

11. J. C. Nauen, G. V. Lauder, Quantification of the wake of rainbow trout (Oncorhynchus mykiss) using three-dimensional stereoscopic digital particle image velocimetry. J. Exp. Biol. 205, 3271–3279 (2002).

12. U.K. Müller, B.L.E. Van Den Heuvel, E.J. Stamhuis, J.J. Videler, Fish foot prints: morphology and energetics of the wake behind a continuously swimming mullet (Chelon labrosus Risso). The J. Exp. Biol. 200, 2893–2906 (1997).

13. E. D. Tytell, “Do trout swim better than eels? Challenges for estimating performance based on the wake of self-propelled bodies” in Animal Locomotion, G. K. Taylor, M. S. Triantafyllou, C. Tropea, Eds. (Springer Berlin Heidelberg, 2010), pp. 63–74.

14. R. Blickhan, C. Krick, D. Zehren, W. Nachtigall, T. Breithaupt, Generation of a vortex chain in the wake of a Subundulatory swimmer. Naturwissenschaften 79, 220–221 (1992).

15. J. C. Liao, D. N. Beal, G. V. Lauder, M. S. Triantafyllou, Fish exploiting vortices decrease muscle activity. Science 302, 1566–1569 (2003).

16. M. Taguchi, J. C. Liao, Rainbow trout consume less oxygen in turbulence: the energetics of swimming behaviors at different speeds. J. Exp. Biol. 214, 1428–1436 (2011).

17. A. P. Maertens, A. Gao, M. S. Triantafyllou, Optimal undulatory swimming for a single fish-like body and for a pair of interacting swimmers. J. Fluid Mech. 813, 301–345 (2017).

18. B. M. Boschitsch, P. A. Dewey, A. J. Smits, Propulsive performance of unsteady tandem hydrofoils in an in-line configuration. Phys. Fluids 26, 051901 (2014).

19. M. Kurt, K. W. Moored, Flow interactions of two- and three-dimensional networked bio-inspired control elements in an in-line arrangement. Bioinspir. Biomim. 13, 045002 (2018).

20. R. L. McLaughlin, D. L. G. Noakes, Going against the flow: an examination of the propulsive movements made by young brook trout in streams. Can. J. Fish. Aquat. Sci. 55, 853–860 (1998).

21. S. W. Shuler, R. B. Nehring, K. D. Fausch, Diel Habitat Selection by Brown Trout in the Rio Grande River, Colorado, after Placement of Boulder Structures. N. Am. J. Fish. Manage. 14, 99–111 (1994).

22. J. M. Anderson, K. Streitlien, D. S. Barrett, M. S. Triantafyllou, Oscillating foils of high propulsive efficiency. J. Fluid Mech. 360, 41–72 (1998).

23. J. H. J. Buchholz, A. J. Smits, On the evolution of the wake structure produced by a low-aspect-ratio pitching panel. J. Fluid Mech. 564, 433–443 (2005).

24. M. Saadat, et al., On the rules for aquatic locomotion. Phys. Rev. Fluids 2, 083102 (2017).

25. K. N. Lucas, G. V. Lauder, E. D. Tytell, Airfoil-like mechanics generate thrust on the anterior body of swimming fishes. Proc. Natl. Acad. Sci. U. S. A. 117, 10585–10592 (2020).

26. K. T. Du Clos, et al., Thrust generation during steady swimming and acceleration from rest in anguilliform swimmers. J. Exp. Biol. 222 (2019).

27. J. O. Dabiri, S. Bose, B. J. Gemmell, S. P. Colin, J. H. Costello, An algorithm to estimate unsteady and quasi-steady pressure fields from velocity field measurements. J. Exp. Biol. 217, 331–336 (2014).

28. S. Verma, G. Novati, P. Koumoutsakos, Efficient collective swimming by harnessing vortices through deep reinforcement learning. Proceedings of the National Academy of Sciences 115, 5849–5854 (2018).

29. D. N. Beal, F. S. Hover, M. S. Triantafyllou, J. C. Liao, G. V. Lauder, Passive propulsion in vortex wakes. J. Fluid Mech. 549, 385–402 (2006).

30. V. D. Santo, et al., Convergence of undulatory swimming kinematics across a diversity of fishes. Proceedings of the National Academy of Sciences 118 (2021).

31. M. F. Steinhausen, J. F. Steffensen, N. G. Andersen, Tail beat frequency as a predictor of swimming speed and oxygen consumption of saithe (Pollachius virens) and whiting (Merlangius merlangus) during forced swimming. Mar. Biol. 148, 197–204 (2005).

32. J. Ohlberger, G. Staaks, F. Hölker, Estimating the active metabolic rate (AMR) in fish based on tail beat frequency (TBF) and body mass. J. Exp. Zool. A Ecol. Genet. Physiol. 307, 296–300 (2007).

33. L. Li, et al., Vortex phase matching as a strategy for schooling in robots and in fish. Nat. Commun. 11, 5408 (2020).

34. I. Borazjani, F. Sotiropoulos, Numerical investigation of the hydrodynamics of carangiform swimming in the transitional and inertial flow regimes. J. Exp. Biol. 211, 1541–1558 (2008).

35. G. Liu, H. Dong, “Effects of Tail Geometries on the Performance and Wake Pattern in Flapping Propulsion” in Proceedings of the ASME 2016 Fluids Engineering Division Summer Meeting. https:/doi.org/10.1115/FEDSM2016-7691. (2016).

36. Domel, et al., Shark skin-inspired designs that improve aerodynamic performance. J. R. Soc. Interface 15, 20170828, (2018).

37. R. Thandiackal, C. H. White, H. Bart-Smith, G. V. Lauder, Tuna robotics: hydrodynamics of rapid linear accelerations. Proc. Biol. Sci. 288, 20202726 (2021).

38. J. Zhu, et al., Tuna robotics: A high-frequency experimental platform exploring the performance space of swimming fishes. Sci Robot 4 (2019).

39. R. Thandiackal, et al., Emergence of robust self-organized undulatory swimming based on local hydrodynamic force sensing. Sci Robot 6 (2021).

40. R. Thandiackal, G. V. Lauder, How zebrafish turn: analysis of pressure force dynamics and mechanical work. J. Exp. Biol. 223, jeb223230 (2020).

